# Digital holo-tomographic 3D maps of COVID-19 microclots in blood to assess disease severity

**DOI:** 10.1101/2023.09.12.557318

**Authors:** Talia Bergaglio, Olena Synhaivska, Peter Niraj Nirmalraj

**Affiliations:** Transport at Nanoscale Interfaces Laboratory, Swiss Federal Laboratories for Materials Science and Technology, Dübendorf CH-8600, Switzerland; Graduate School for Cellular and Biomedical Sciences, University of Bern, Bern CH-3012, Switzerland

**Keywords:** microclots, plasma, COVID-19, long COVID, label-free imaging

## Abstract

The coronavirus disease 2019 (COVID-19) has impacted health globally. Cumulative evidence points to long-term effects of COVID-19 such as cardiovascular and cognitive disorders diagnosed in patients even after the recovery period. In particular, micrometer-sized blood clots and hyperactivated platelets have been identified as potential indicators of long COVID. Here we resolve individual microclot structures in platelet-rich plasma of donors with different subphenotypes of COVID-19 in a label-free manner, using 3D digital holo-tomographic microscopy (DHTM). Based on 3D refractive index (RI) tomograms, the size, dry mass, and prevalence of microclot composites were quantified and then parametrically differentiated from fibrin-rich microclots and platelet aggregates in the plasma of COVID-19 donors. Importantly, fewer microclots and platelet aggregates were detected in the plasma of healthy controls when compared to COVID-19 donors. Our work highlights the utility of integrating DHTM in clinical settings that may allow the detection of individuals at risk of developing microvascular thrombotic disorders and for monitoring the efficiency of prescribed treatments by screening plasma samples.

## Main

Coronavirus disease 2019 (COVID-19) is a complex disorder caused by infection with severe acute respiratory syndrome coronavirus 2 (SARS-CoV-2) and affects many bodily functions, including excessive immunological response, autoimmunity, and endothelial dysfunction, that have been implicated in the risk of thrombotic events and coagulopathies (*1-7*). While the acute phase of COVID-19 primarily manifests as a respiratory illness, with a wide range of symptoms including fever, headache, cough, fatigue, muscle pain, and shortness of breath, more severe clinical symptoms, such as systemic inflammatory response syndrome (SIRS), acute respiratory disease syndrome (ARDS), neurological and cardiovascular complications, have been associated with processes underlying coagulopathies and endotheliopathies (*4, 6, 8*). Currently, it is estimated that approximately 30% of individuals with COVID-19 continue to suffer from a variety of different symptoms involving specific or multiple organ systems, with neurological, neuropsychiatric, and cardiorespiratory clinical presentations (*8, 9*), a condition known as post-acute sequelae of COVID (PASC) or long COVID (*8, 10, 11*). Fatigue, cough, dyspnea, chest pain, headache, dizziness, cognitive impairment known as ‘brain fog’, and long-term smell and taste dysfunction are the most commonly reported symptoms in individuals suffering from long COVID (*9, 10, 12*). The World Health Organization defines long COVID as a condition in which individuals with a probable or confirmed infection with SARS-CoV-2 still experience COVID-19-related symptoms usually three months post-infection and lasting for at least two months, with no alternative diagnosis (*13*). Endotheliopathy, coagulopathy, and thrombosis are also now established complications of COVID-19 and can persist in individuals diagnosed with long COVID, contributing to the pathogenesis and clinical manifestations of these conditions (*1, 14-16*).

The pathogenic mechanisms underlying coagulopathy in COVID-19 and long COVID are complex and multifactorial, including a patient’s pre-existing conditions (*17, 18*). The presence of SARS-CoV-2 has been detected within endothelial cells, suggesting a direct viral effect on the vascular system (*19, 20*). Additionally, the dysregulated immune response triggered by the viral infection can lead to excessive inflammation and cytokine release, further promoting a pro-thrombotic state (*21*). More specifically, COVID-19-related coagulopathy was associated with elevated levels of von Willebrand Factor (VWF), D-dimer, fibrinogen, and markers of platelet activation, as well as damaged red blood cells (RBCs) and reduced fibrinolysis, reflecting ongoing coagulation activation (*3, 22, 23*). In long COVID, persistent coagulation abnormalities may contribute to the chronic and debilitating symptoms experienced by affected individuals (*1, 8, 24*). The formation of microclots, or microthrombi, within the pulmonary vasculature has been observed, contributing to respiratory compromise (*5, 25, 26*). The mechanisms underlying these persistent coagulation disturbances are not yet fully understood, but they may involve a combination of residual inflammation, immune dysregulation, and vascular damage (*2, 5, 7*).

The incidence of microvascular injury in COVID-19 patients has been detected in the lungs, brain, heart, and other organs during autopsy, using magnetic resonance imaging (MRI) and upon histological examination (*24, 27-29*). Previously, Pretorius and co-workers (*26*) reported on the presence of microclots in COVID-19 platelet-poor plasma (PPP), using thioflavin T (ThT) staining and fluorescent microscopy. These microclots, ranging in size between 1 and 200 μm, were determined to be amyloidogenic, indicated by the positive ThT signal, suggesting a more anomalous clot structure characterized by extensive ß-sheet structures and presenting resistance to fibrinolysis (*4, 5, 20, 30, 31*). Due to their size, microclots are known to block micro-capillaries and prevent efficient oxygen transport through the blood circulation, resulting in a cascade of hemolytic events responsible for part if, not most of, the symptomatology in both acute and long COVID (*5, 21, 32*).

Scanning electron microscopy and microfluidic-based studies have also provided insights into the coagulation mechanisms in COVID-19 and long COVID compared to healthy plasma by inducing blood clot formation with/without thrombin and SARS-CoV-2 spike protein (*20, 33*). Denser structures with thicker fibrils in the range of ∼50-150 nm in diameter were observed, in addition to increased clot rigidity and platelet hyperactivation in the COVID-19 samples. Proteomics analysis of microclots revealed the presence of fibrin in amyloid form, platelets, and additional plasma proteins, including VWF, complement factors, C-reactive protein, α2-antiplasmin and the SARS-CoV-2 virus, which may induce the production of different prothrombotic autoantibodies (*3, 5, 33-35*).

To date, evidence for the presence of microclots in the plasma of COVID-19 and long COVID donors has been mainly obtained using *in vitro* ThT staining and fluorescent microscopy (*4, 9, 26*). Currently available diagnostic methods for the assessment of coagulopathies include qualitative assessment using ThT labeling-based fluorescent microscopy, or using semi-quantitative Thromboelastography (TEG) and D-dimer analysis (*3-5, 36*). Importantly, the presence of high levels of D-dimer can serve as an indicator for the production of blood clots and subsequent adequate fibrinolysis but does not sufficiently reflect the nature, composition, and number of the residual microclots in circulation (*5*). Hence, it would be highly desirable to devise a methodology that would allow for direct determination of microclot morphology, composition, and prevalence in blood in a label-free manner compatible with operating even in resource-limited clinical settings. To address this pending need, we have developed an imaging methodology based on digital holo-tomographic microscopy (DHTM) to three-dimensionally map microclot structure and composition in plasma samples from donors diagnosed with COVID-19, with high spatial resolution and in a label-free manner.

First, the refractive index (RI) maps of synthetically prepared blood clot fragments in aqueous solution from healthy individuals and donors with COVID-19 were registered using DHTM (see Methods section for details on clot synthesis procedure). The structural information and blood clot composition were quantified by segmenting and digitally staining the 3D RI tomograms, to identify differences in size and composition between blood clot types. Raman spectroscopy measurements were also conducted to further characterize the blood clot structure and composition. Next, we extended the imaging and analysis protocols to directly examine the platelet-rich plasma (PRP) of individuals with different COVID-19 subphenotypes, including COVID-19 positive and recovered. Microclots with varying structure and composition were detected in plasma samples from all the donors with COVID-19, regardless of symptom severity. Microclots were classified based on their composition as either microclot composites, fibrin-rich microclots or platelet aggregates. To understand whether microclot structure and composition would differ between COVID-19 subphenotypes, we quantified the overall size distribution and prevalence of microclot subtypes in PRP from COVID-19 compared to healthy donors. Our findings highlight that label-free high-throughput microscopy can be used as a point-of-care technique to visualize and quantify the presence of microclots directly in plasma without the need for complicated sample preparation techniques. Knowledge gained from such microclot analytics could be relevant for developing effective treatment strategies to prevent and manage thrombotic complications in COVID-19 and long COVID patients, thereby improving patient outcomes.

## Results

### Label-free digital holo-tomographic microscopy of synthetically prepared blood clots

Previously, we have shown that DHTM can be used to visualize and quantify the morphometry of red blood cells as they interact with ibuprofen drugs in a non-invasive and label-free manner, with high spatial resolution (*37*). In the current study, we first extended the use of DHTM (Supplementary Figure 1) to resolve the structure and composition of synthetically prepared blood clot fragments in aqueous solution (Fig. 1). Blood from a healthy and a convalescent COVID-19 donor was collected and allowed to clot before imaging with DHTM (Methods). Details on the demographic information of the donors are provided in Supplementary Table 1. Figure 1a shows the 3D refractive index (RI) tomogram of synthetically prepared blood clot fragments from a healthy donor, revealing differences in the structural composition based on variations in RI values. From the corresponding segmented RI tomogram (Fig. 1b), micrometer-sized blood clot fragments were digitally stained based on the RI values (Fig. 1c) and a color-coded label was assigned to discriminate between different structural components, including fibrin strands and non-fibrin structures. The same RI classification was applied to spatially well-resolved blood clot fragments obtained from a COVID-19 donor (Fig. 1d), revealing similar structural components depicting a net-like fibrin network with smaller and circular non-fibrin structures.

**Fig. 1:**
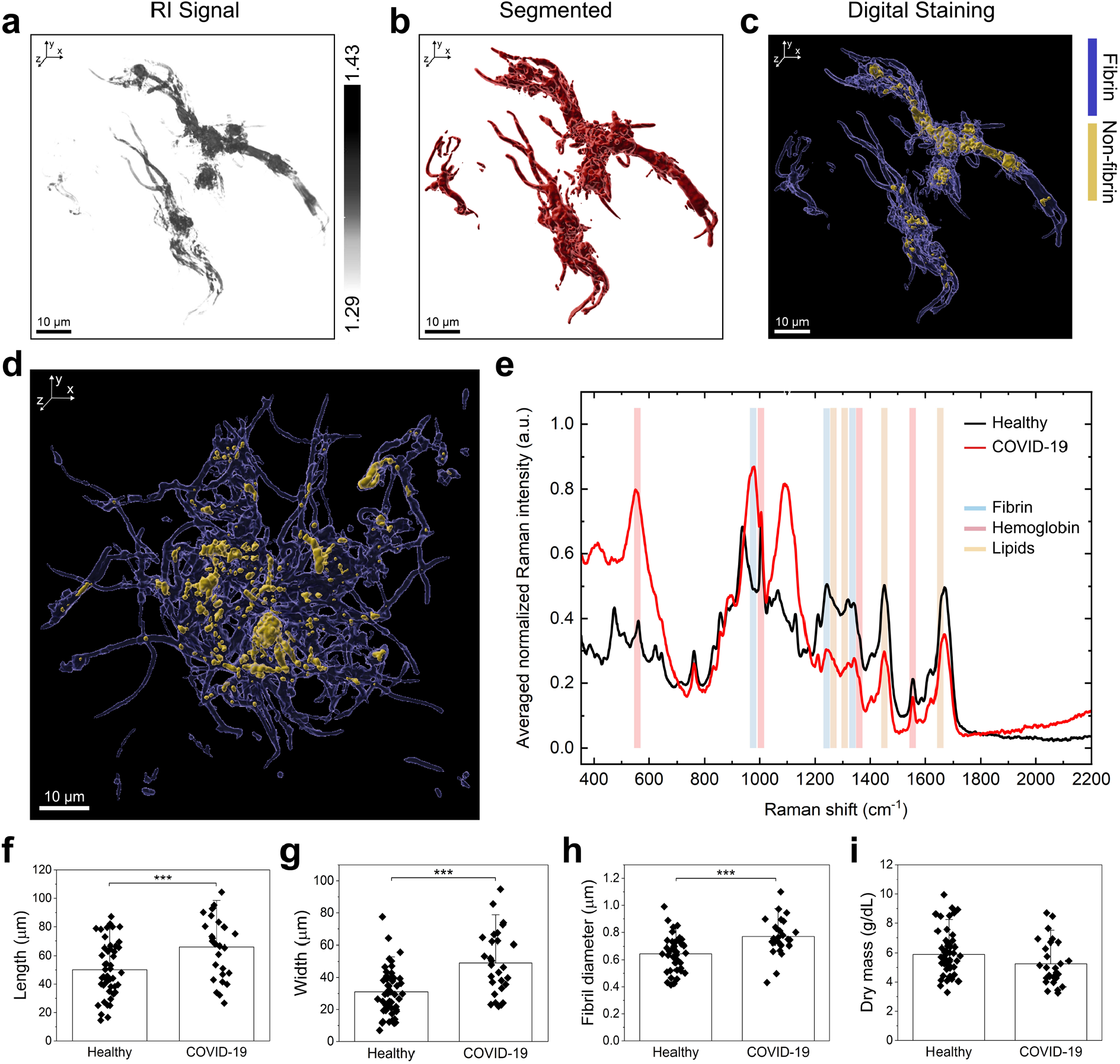
Structural and spectrochemical analysis of synthetically prepared fixed blood clots in aqueous solution. **a**, Refractive index (RI) tomogram of a blood clot fragment obtained from a healthy donor. **b**, Corresponding segmented RI tomogram and **c**, digital RI staining for fibrin (blue) and non-fibrin structures (yellow). **d**, Digital RI staining for fibrin (blue) and non-fibrin structures (yellow) of a blood clot obtained from a donor with COVID-19. **e**, Averaged spectral signatures of blood clots obtained from a healthy donor (black line) and a donor with COVID-19 (red line). Quantification of **f**, length, **g**, width, **h**, fibrin fibril diameter and **i**, dry mass of blood clots from a healthy donor and a donor with COVID-19.

To further understand the compositional differences within a microclot structure, we performed Raman spectroscopy on the same samples (Methods) used previously for DHTM measurements based on which data are shown in Fig. 1a and d were obtained. Figure 1e shows the Raman spectra collected from microclots of healthy (black spectrum) and COVID-19 (red spectrum) donors. Spectrochemical analysis revealed the presence of signal peaks in the spectral regions associated with the presence of fibrin (976, 1248, and 1342 cm^-1^) (*38*), hemoglobin (567, 1000, 1368, 1542, and 1575 cm^-1^) (*38, 39*), and lipids (1266, 1300, 1444 and 1655 cm^-1^) (*40*). The averaged spectral signature of blood clots from a donor with COVID-19 presented spectral similarities to the healthy blood clots in the hemoglobin and lipid regions. In the hemoglobin region, an increase in the intensity of the peak at 561 cm^-1^ was detected in the COVID-19 blood clots, indicative of stretching of the Fe-O_2_ bond (*39*). In contrast, a decrease in the intensity of the peak at 1360 cm^-1^ was observed in the COVID-19 compared to the healthy blood clots. In the lipid regions, a decrease in the intensity of the peaks at 1266 cm^-1^ and 1655 cm^-1^ as well as the peaks at 1305 cm^-1^ and 1450 cm^-1^ was detected in the COVID-19 blood clot samples. These peaks are associated with unsaturated and saturated lipids, respectively (*40*). In the fibrin regions, a maximum peak at 976 cm^-1^ was detected in the COVID-19 but not in the healthy blood clots. Here, the presence of a peak at 937 cm^-1^ may be indicative of a shift of the fibrin band in the healthy blood clots. In contrast, both blood clot samples presented a maximum peak in a secondary fibrin region at 1342 cm^-1^, with a lower signal intensity in the COVID-19 blood clots. Such differences in the spectra between the healthy and COVID-19 blood clot samples may be indicative of changes occurring in the overall structure and composition of blood clots.

The structural parameters of healthy and COVID-19 blood clot fragments were extracted and quantified, including the overall size, the fibrin fibril diameter, and the dry mass. A two-sample t-test revealed a significant increase in clot length (healthy: 50.1 ± 18.9 μm; COVID-19: 66.2 ± 21.5 μm) (Fig. 1f) and width (healthy: 31.0 ± 14.5 μm; COVID-19: 49.1 ± 19.8 μm) (Fig. 1g) in COVID-19 compared to healthy blood clots, suggesting an overall difference in the size of the clot fragments. In addition, the diameter of the fibrin fibrils was significantly larger in the COVID-19 compared to the healthy samples (healthy: 0.64 ± 0.13 μm; COVID-19: 0.77 ± 0.14 μm) (Fig. 1h). Finally, the dry mass was obtained from the RI measurements (Methods). No significant difference was detected between the healthy (5.90 ± 1.57 g/dL) and COVID-19 (5.26 ± 1.5 g/dL) samples (Fig. 1i), indicating a comparable structural composition of the analyzed blood clot fragments.

### Structural analysis of microclots in COVID-19 platelet-rich plasma

The described DHTM imaging and analysis protocol was employed to assess the presence of microclots in platelet-rich plasma (PRP) of five donors with COVID-19 (Supplementary Table 2). Different COVID-19 subphenotypes were identified, depending on the presence (positive or recovered) and severity of symptoms (mild, moderate, asymptomatic) at the time of blood collection. Microclots of varying sizes were observed during DHTM measurements in plasma from all five donors. The detected microclots were classified based on their composition: (i) microclot composites, defined as micrometer-size clot structures with mixed composition of plasma proteins (*34*); (ii) fibrin–rich microclots, defined as micrometer-size clot structures with prevalent fibrin fibrils and (iii) platelet aggregates, defined as aggregated platelets of different sizes depending on the number of platelets (<10 platelets, 10-50 platelets and >50 platelets). Microclots were found in all the analyzed COVID-19 plasma samples, regardless of age, symptomatology, and IgG and IgM antibody levels. Figure 2 shows the results of the DHTM-based analysis of plasma collected from a 27-year-old female donor (donor identified as sample PRP-CVDP2) with a COVID-19 positive and mild symptomatology subphenotype (Methods). The 3D RI tomograms show highly aggregated platelets and the presence of a fibrin-rich microclot of moderate size (∼35 μm) (Fig. 2a,c). The microclot structures were segmented and classified based on the microclot type and platelet aggregate size (Fig. 2b,d). To morphologically characterize the microclots, the size, and the dry mass were extracted and quantified from the RI tomograms (Methods). Fibrin-rich microclots were bigger (mean = 23.2 ± 7.9 μm) compared to microclot composites (mean = 16.5 ± 5.6 μm), likely due to the length of the fibrin strands in a network format (Fig. 2e). In contrast, the dry mass did not differ between the microclot structures (fibrin-rich microclots = 14.2 ± 0.7 g/dL, microclot composite = 14.9 ± 0.7 g/dL) (Fig. 2f). The size of platelet aggregates differed markedly based on the number of platelets (<10 platelets = 2.6 ± 1.3 μm, 10-50 platelets = 13.8 ± 8.2 μm, >50 platelets = 26.8 ± 9.1 μm), with platelet aggregates of >50 platelets reaching sizes of up to 45 μm (Fig. 2g). No major differences were observed in the dry mass of the platelet aggregates (<10 platelets = 15.3 ± 1.3 g/dL, 10-50 platelets = 15.7 ± 1.0 g/dL, >50 platelets = 15.7 ± 0.7 g/dL) (Fig. 2h), likely because these microclot structures are composed of mostly platelets. A total of ∼4 microclot composite structures, ∼5 fibrin-rich microclots, and ∼1570 platelet aggregates were observed in 75 μL of plasma (Fig. 2i), indicating extensive platelet pathology characterized by aggregated platelets in plasma from donor PRP-CVDP2.

**Fig. 2:**
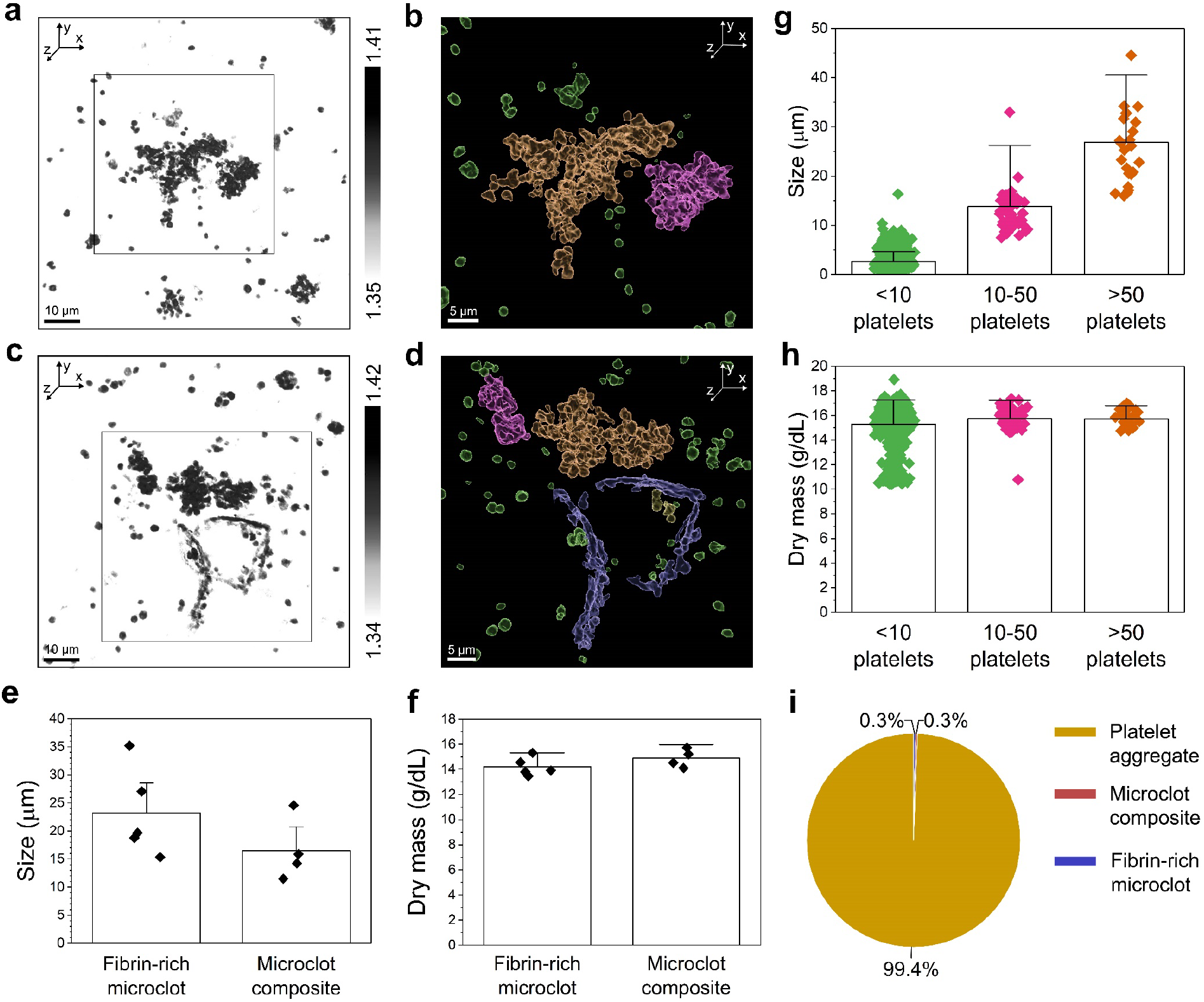
Structural analysis of microclots in platelet-rich plasma (PRP) of a COVID-19 positive donor with mild symptomatology (PRP-CVDP2). **a**, Refractive index (RI) tomogram of platelet aggregates in PRP. **b**, Corresponding segmented RI tomogram of the platelet aggregates of different sizes from the inset in **a. c**, RI tomogram of a fibrin-rich microclot and platelet aggregates in PRP. **d**, Corresponding segmented RI tomogram of the fibrin-rich microclot and the platelet aggregates of different sizes from the inset in **c. e**,**f**, Size and dry mass of fibrin-rich microclots and microclot composite structures in PRP. **g**,**h**, Size and dry mass of platelet aggregates in PRP. Error bars represent the standard deviation from the mean. **i**, Pie chart showing the microclot composition in PRP of donor PRP-CVDP2.

Next, we analyzed the plasma of a 26-year-old female donor (donor identified as sample PRP-CVDP1) with a COVID-19 positive and moderate symptomatology subphenotype (Fig. 3 and Methods). A total of ∼20 microclots were detected in 75 μL of PRP, classified as either microclot composite (Fig. 3a) or fibrin-rich microclots (Fig. 3c,e). The segmented and classified 3D RI tomograms show the spatially magnified structure of a microclot composite (Fig. 3b) and reveal the presence of single fibrin fibrils in the plasma of this individual (Fig. 3d,f). A clear size difference was observed between the two types of microclot structures (Fig. 3g). Similar to the PRP-CVDP2, the measured fibrin-rich microclots appeared larger (mean = 24.0 ± 6.7 μm) compared to the microclot composites (mean = 8.1 ± 8.0 μm). Interestingly, the dry mass did not differ between microclot structures (fibrin-rich microclots = 12.9 ± 0.5 g/dL, microclot composite = 13.4 ± 1.9 g/dL), except two microclot composites (Fig. 3h). Overall, the described COVID-19 subphenotype was characterized by prevalent microclot composite structures and a few fibrin-rich microclots (Fig. 3i). Platelet clumping was not observed in plasma from donor PRP-CVDP1.

**Fig. 3:**
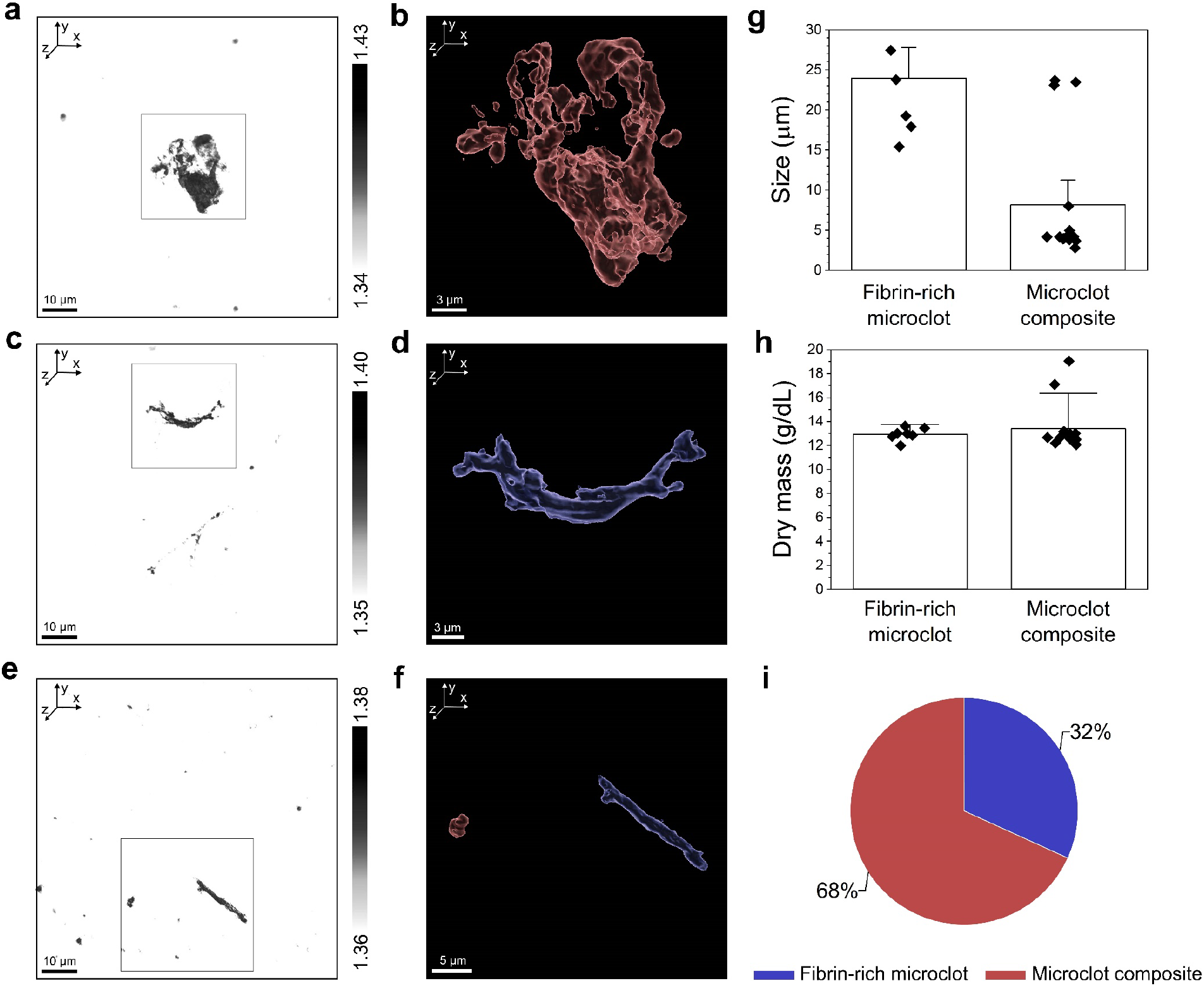
Structural analysis of microclots in platelet-rich plasma (PRP) of a COVID-19 positive donor with moderate symptomatology (PRP-CVDP1). **a**, Refractive index (RI) tomogram of a microclot composite in PRP. **b**, Corresponding segmented RI tomogram of the microclot composite from the inset in **a. c**, RI tomogram of a fibrin-rich microclot in PRP. **d**, Corresponding segmented RI tomogram of the fibrin-rich microclot from the inset in **c. e**, RI tomogram of a microclot composite and fibrin-rich microclots in PRP. **f**, Corresponding segmented RI tomogram of a microclot composite and the fibrin-rich microclot from the inset in **e. g**,**h**, Size and dry mass of fibrin-rich microclots and microclot composite structures in PRP. Error bars represent the standard deviation from the mean. **i**, Pie chart showing the microclot composition in PRP of donor PRP-CVDP1.

Finally, we report the analysis of plasma collected from a 23-year-old female donor (donor identified as sample PRP-CVDR2) with a recovered COVID-19 and mild symptomatology subphenotype (Fig. 4 and Methods). The presence of microclots was detected in 75 μL of PRP, despite this donor being classified as COVID-19 recovered and no longer experiencing symptoms at the time of blood collection. Microclot composites (n = 16) presented varying morphologies, including more dense structures with fibrin fibrils (Fig. 4a,b) and net-like structures intertwined with platelets (Fig. 4c,d,e,f). Platelet clumping was also observed (n = 83) (Fig. 4g,h). The size of fibrin-rich microclots and microclot composites did not significantly differ (fibrin-rich microclots = 19.0 ± 1.6 μm, microclot composites = 20.1 ± 19.1 μm) as shown in Fig. 4i. However, a few large microclot composite structures were detected, measuring up to ∼80 μm in size. Similarly, the dry mass was consistent between the microclot structures (fibrin-rich microclots = 14.3 ± 0.6 g/dL, microclot composite = 14.1 ± 1.3 g/dL) (Fig. 4j). Platelet aggregates varied in size depending on the number of platelets (<10 platelets = 3.5 ± 1.9 μm, 10-50 platelets = 15.5 ± 3.3 μm, >50 platelets = 37.9 ± 7.2 μm), reaching sizes of up to ∼50 μm (Fig. 4k).

**Fig. 4:**
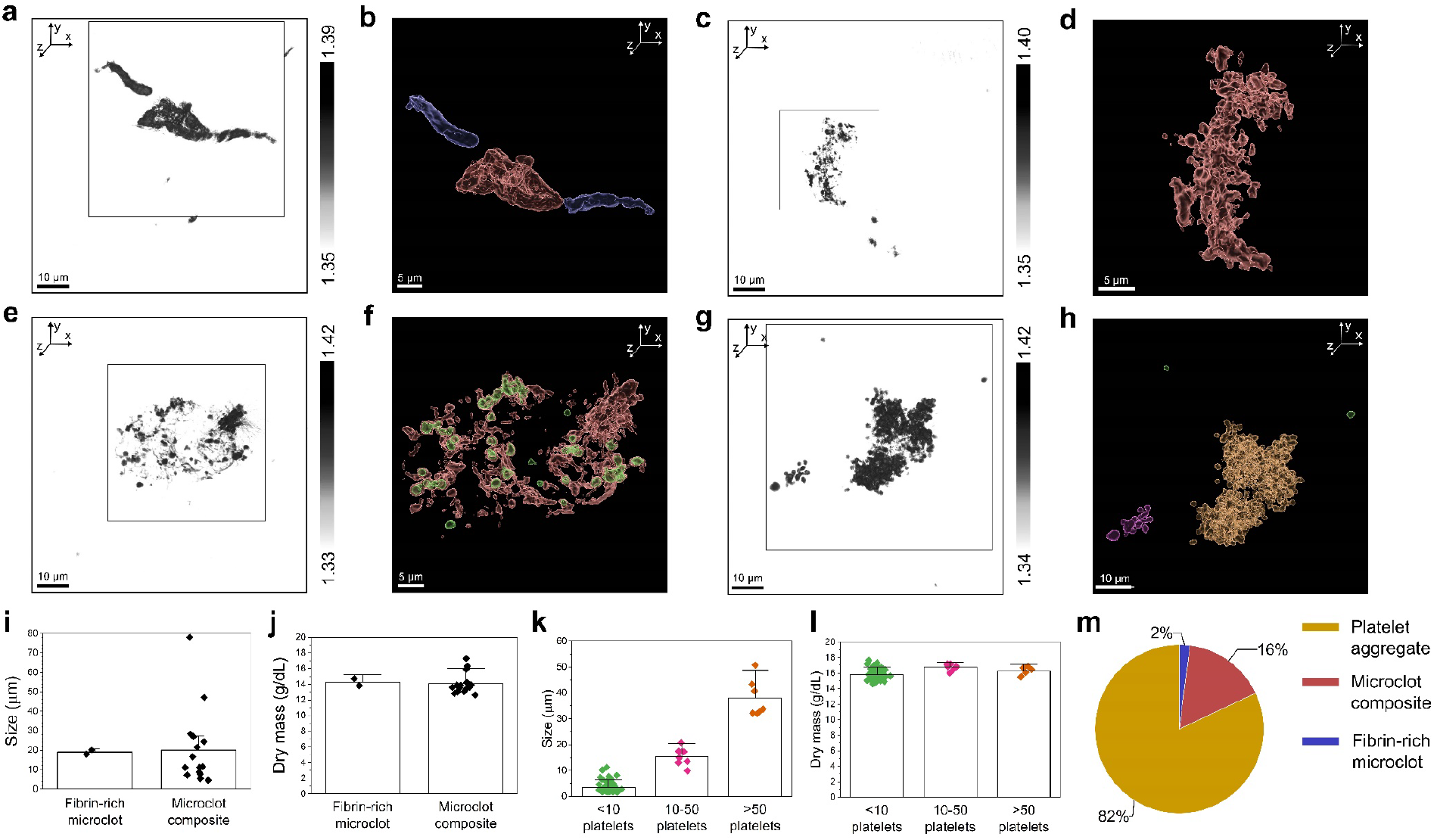
Structural analysis of microclots in platelet-rich plasma (PRP) of a COVID-19 recovered donor with mild symptomatology (PRP-CVDR2). **a**, Refractive index (RI) tomogram of a microclot composite and fibrin-rich microclots in PRP. **b**, Corresponding segmented RI tomogram of the microclot composite and fibrin-rich microclots from the inset in **a. c**, RI tomogram of a microclot composite in PRP. **d**, Corresponding segmented RI tomogram of the microclot composite from the inset in **c. e**, RI tomogram of a microclot composite with platelet aggregates in PRP. **f**, Corresponding segmented RI tomogram of the microclot composite with platelet aggregates from the inset in **e. g**, RI tomogram of platelet aggregates in PRP. **h**, Corresponding segmented RI tomogram of the platelet aggregates of different sizes from the inset in **g**. i,**j**, Size and dry mass of fibrin-rich microclots and microclot composite structures in PRP. **k**,**l**, Size and dry mass of platelet aggregates in PRP. Error bars represent the standard deviation from the mean. **m**, Pie chart showing the microclot composition in PRP of donor PRP-CVDR2.

Instead, the dry mass did not differ between platelet aggregate structures (<10 platelets = 15.8 ± 0.6 g/dL, 10-50 platelets = 16.8 ± 0.4 g/dL, >50 platelets = 16.3 ± 0.6 g/dL) (Fig. 4l). Overall, this diverse COVID-19 subphenotype presented a mixed composition of microclot structures, with platelet clumping representing the most prevalent feature (Fig. 4m). The structural analysis of the remaining COVID-19 subphenotypes, including a COVID-19 positive female donor with no symptomatology (PRP-CVD3) and a COVID-19 recovered female donor with moderate symptomatology (PRP-CVDR1), and of the healthy controls are provided in the Supplementary figure 2, 3 and 4, respectively. Interestingly, microclots were detected in plasma from PRP-CVD3 donors even in the absence of clinical features (Supplementary Figure 2).

### Quantification of microclot structure and composition in COVID-19 subphenotypes

Following the in-depth description of the DHTM analysis of PRP from the single COVID-19 donors, we present the results on the quantification of microclot structure and composition of all COVID-19 subphenotypes as well as five healthy controls (Fig. 5). Figure 5a shows the prevalence of fibrin-rich microclots and microclot composites in 75 μL of PRP from five healthy donors and five donors with different COVID-19 subphenotypes. Overall, these microclot structures were more prevalent in PRP from all COVID-19 samples compared to the healthy controls (fibrin-rich microclot: 1 ± 2; microclot composite: 2 ± 2). Within the COVID-19 samples, microclot composites were more prevalent compared to fibrin-rich microclots, except for the COVID-19 positive donor with mild symptomatology, which presented an equal number of microclot structures (Fig. 5a). Additionally, a trend for a higher prevalence of fibrin-rich microclots in COVID-19 positive compared to COVID-19 recovered donors can be observed (Fig. 5a). Platelet clumping was also more prevalent in the COVID-19 positive compared to the recovered subphenotypes, although it was not observed in the COVID-19 positive donor with moderate symptomatology (Fig. 5b-e). Aggregated platelets were not detected in any of the healthy samples. In all samples where platelet clumping was detected, platelet aggregates of <10 platelets were mostly prevalent (Fig. 5b-e). Interestingly, larger platelet aggregates composed of >10 platelets were detected in the PRP of both COVID-19 donors (positive and recovered) with mild symptomatology (Fig. 5b-e). However, a correlation between microclot type and prevalence with the analyzed COVID-19 subphenotypes could not be determined.

**Fig. 5:**
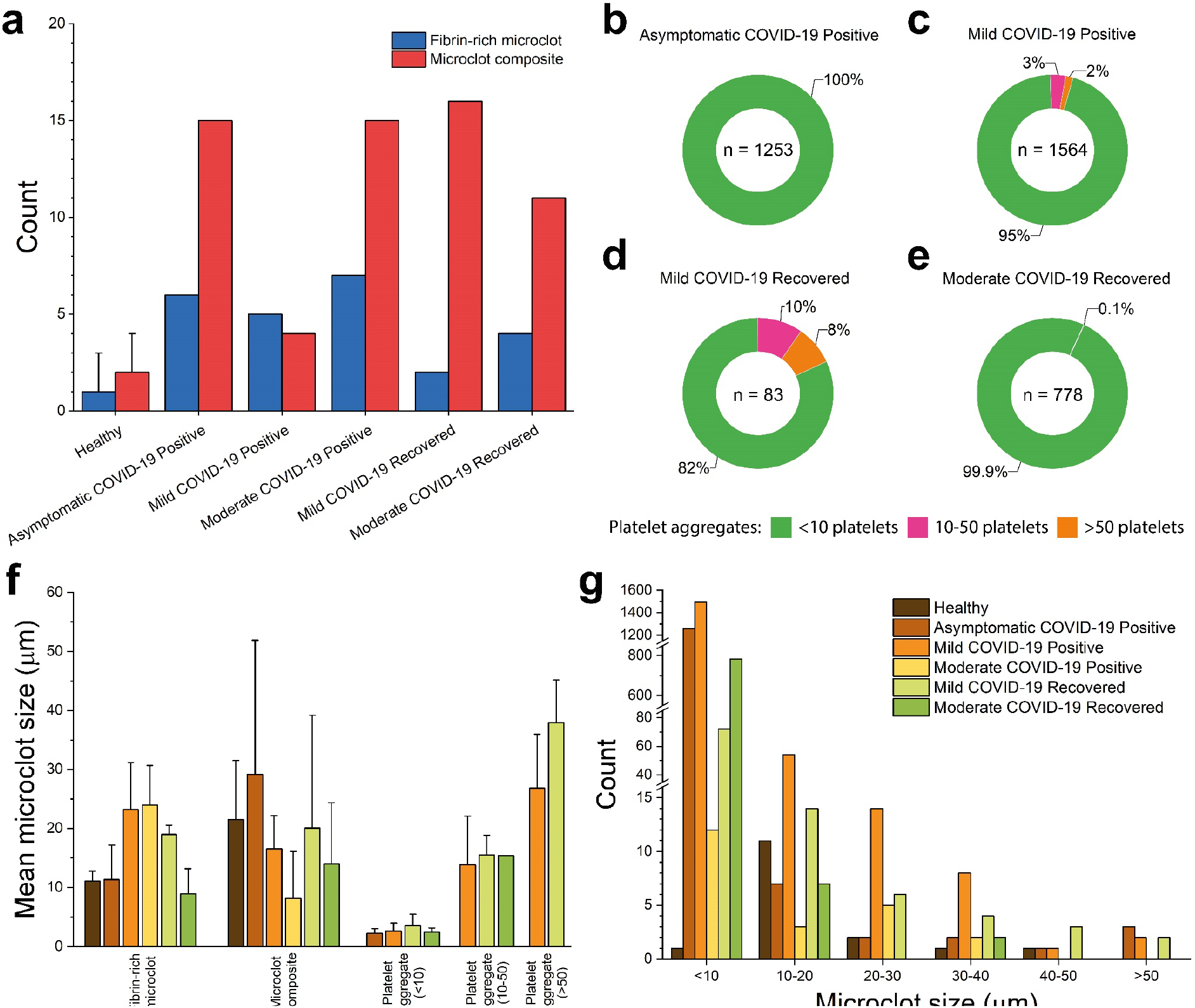
Morphological characterization of microclots in healthy and COVID-19 positive and recovered donors. **a**, Prevalence of fibrin-rich microclots and microclot composites detected in 75 μL of PRP from healthy and COVID-19 positive and recovered donors. **b**, Prevalence of platelet aggregates detected in 75 μL of PRP from a COVID-19 positive donor with no symptomatology, **c**, a COVID-19 positive donor with mild symptomatology, **d**, a COVID-19 recovered donor with mild symptomatology and **e**, a COVID-19 recovered donor with moderate symptomatology. **f**, Mean microclot size plotted against the microclot type and grouped by healthy and COVID-19 subphenotypes. Error bars correspond to the standard deviation. **g**, Prevalence of microclots plotted against the microclot size and grouped by healthy and COVID-19 subphenotypes.

Due to the importance of microclot size in the risk of occlusion of micro-capillaries, the mean microclot size was plotted against the microclot type and grouped by healthy and COVID-19 subphenotypes (Fig. 5f). The size of the detected microclots averaged between 1 and 60 μm. Size variation within fibrin-rich microclots and microclot composites between PRP samples was likely dependent on the length of the fibrin fibrils and on the mixed composition of the microclot composites, respectively (Fig. 5f). The average size of platelet aggregates was comparable between COVID-19 subphenotypes and depended on the number of platelets in each aggregate (Fig. 5f). The prevalence of microclots was also plotted against the microclot size and grouped by healthy and COVID-19 subphenotypes (Fig. 5g). For all COVID-19 subphenotypes, microclots were mostly prevalent in the size range between 1 and 10 μm. Overall, the severity of COVID-19 symptomatology as well as the positive or recovered subphenotype did not seem to correlate with the size of the measured microclots. Moreover, a correlation between microclot prevalence and size and the donor age as well as the IgG and IgM antibody levels was not observed (Supplementary Figures 5 and 6) within the limited number of patient samples analyzed in the present study.

## Discussion

Microclots continue to receive clinical research interest as potential indicators of long-term COVID (*1, 3-5, 8, 9, 12, 15, 20, 26, 36*). In this study, we employed DHTM as an analytical tool to characterize the composition of platelet-rich plasma from donors with COVID-19 subphenotypes. To test our imaging and analysis protocol, we studied synthetically prepared fixed blood clot fragments in an aqueous solution using DHTM. The preliminary step was to identify and obtain the quantitative morphological parameters from 3D RI tomograms of synthetically prepared blood clot fragments. Blood clot structure can be described by parameters such as the fibrin fibril diameter, fibrin length, clot composition, clot density, and clot porosity (*41*). These physical parameters also provide deeper insights into clot stiffness, stability, and degradation dynamics, which play a crucial role in hemolytic processes (*30, 41, 42*). Significant size differences were observed between healthy and COVID-19 blood clot fragments, with regard to the size and fibrin fibril thickness. Although these blood clots were passed through a 30 μm filter in order to fragment them, in general bigger fragments, evidenced by thicker fibrin fibrils, derived from COVID-19 donors suggestive of increased clot rigidity. Clot composition and density, estimated from the dry mass, were similar between the two samples, also supported by the spectrochemical signature from Raman measurements. Importantly, digital staining revealed details of an intricate network of fibrin fibrils with smaller and more circular structures trapped in between the fibrils. Similar to fluorescent markers in immunofluorescence, digital staining allows for specific labeling of multiple structures based on the refractive index values, but without altering the inherent features of the sample (*43*). Although we were able to demonstrate the structural characterization of fixed blood clot fragments in an aqueous solution, we cannot explicitly identify the non-fibrin structures observed within these clot fragments. Taken together, DHTM analysis of micrometer-sized blood clot fragments reveals remarkable details of clot structure and composition in a label-free manner, which can be quantified and further classified into different blood clot types. The morphological parameters that we have identified in the first part of our study were then further extended to directly characterize microclots in plasma.

Next, we extended our DHTM-based imaging protocol to study the composition of platelet-rich plasma from donors with COVID-19 and from healthy donors. Similar to previous studies (*4, 9, 20, 23, 26, 36*), we observed microclots in all COVID-19 plasma samples. However, only fewer microclots were in plasma from healthy donors, suggesting microclotting as a salient feature of COVID-19. One of the most notable findings is the structural diversity of the observed microclots, which we have further classified based on their composition: microclot composites, fibrin-rich microclots and platelet aggregates. Microclot composites were found in all COVID-19 plasma samples, with the highest prevalence in the COVID-19 positive donor with moderate symptomatology. These composites structurally resemble the microclots previously reported by Pretorius and colleagues using fluorescence microscopy (*4, 9, 26, 36*), with a likely mixed composition of fibrin and other proteins (*3, 5, 34*). In some cases, platelets appear to be trapped in between the microclot composite structures. The structural variation in microclot composites observed throughout the COVID-19 plasma samples evidences the presence of a mixed population of fibrin structures and proteins. Microclots were also detected in the form of long fibrin networks. Single fibrin fibrils were also observed, which may stem from clot degradation and fibrinolysis (*4, 20, 29, 42*). Consistent with previous studies (*4, 9, 36*), platelet activation and persistent platelet clumping were prevalent features of platelet pathology in all COVID-19 plasma samples, with the exception of the COVID-19 positive donor with moderate symptomatology. Interestingly, the plasma of this donor was characterized by microclot composites and fibrin-rich microclots only, albeit presenting the most severe clinical presentation and high levels of IgG antibodies. Importantly, the size of the measured microclots in COVID-19 plasma samples using DHTM could be of potential clinical interest in regards to their ability to occlude microvasculature and thus impede oxygen transport (*5*). Microclots formed in the vascular system are likely to explain the heterogeneous symptomatology and multi-organ dysfunction observed in COVID-19 and long COVID patients (*14, 44*).

On the basis of our DHTM analysis, a correlation between microclot type, size and prevalence and COVID-19 subphenotypes could not be established in the small analyzed cohort. However, it is possible that many more pre-existing individual risk factors, aside from SARS-CoV-2 infection status (COVID-19 positive or recovered) and symptomatology, may contribute to the presence and to the extent of COVID-19-related microvascular injury, including a history of smoking, vitamin D deficiency, exposure to air pollutants and the presence of other pathologies such as cancer or diabetes mellitus, for which we could not account for in our study (*1, 5, 6, 26*). The identification of comorbidities is particularly relevant in assessing an individual’s risk of developing long COVID (*9, 16, 18*). It is important to note that the symptomatology reported for the analyzed cohort (e.g. headache, fever) may not directly reflect the coagulopathy status of the donors upon SARS-CoV-2 infection, which in turn would explain why we did not observe a distinct correlation between microclot prevalence and size with COVID-19 subphenotypes. The role of potential risk factors associated with the presence of microclots and the wide spectrum of clinical manifestations of both acute and long COVID may only be unraveled with a larger cohort(*45*). Future studies aimed at validating our DHTM methodology will require a larger sample size to be analyzed from both acute COVID-19 and long COVID conditions and sample stratification needs to be included in the study for identification of different comorbidities and risk factors, as well as differentiation between SARS-CoV-2 variants.

In conclusion, the label-free DHTM-based approach demonstrated in our study could serve as a powerful tool for screening microclots in plasma. Information obtained in a label-free manner from such an approach, with high spatial resolution, could provide deeper insights into understanding the role of microclots in health and disease. We anticipate that such an analytical technique at a single microclot level, compatible with operation under standard laboratory conditions, could be easily integrated with existing clinical pipeline for early detection of microvascular thrombotic and related disorders.

## Methods

### Patient characteristics

In this study, we present data from healthy donors and donors who were either recovering or had recovered from coronavirus 2 (SARS-coV-2) infection, i.e., COVID-19 donors, upon blood collection. Fixed blood clots fragments in aqueous solution were prepared from fresh human blood collected from one healthy donor and one COVID-19 convalescent donor in the United States (Zenbio). In contrast, platelet-rich plasma (PRP) was collected from five healthy donors in the United Kingdom (BioIVT) and five adult COVID-19 convalescent donors in the United States (BioIVT). Three COVID-19 donors were classified as COVID positive as they tested positive for the SARS-coV-2 serology test (IgG/IgM positive, IgG positive/IgM negative) and they were symptomatic at the time of blood collection. Blood samples were collected within one month from the diagnosis of SARS-coV-2 infection. In contrast, two COVID-19 donors were classified as COVID recovered as they were no longer symptomatic at the time of blood collection and presented with either a positive or negative serology test (IgG positive/IgM negative and IgG/IgM negative). Blood samples were collected between two to three months after COVID-19 diagnosis. The type and severity of symptoms in COVID-19 patients were heterogeneous. Within the COVID positive donor group, one was asymptomatic, one presented mild symptoms (body or muscle aches, chills, and headache) and one had moderate symptoms (body or muscle aches, chest pain, chills, confusion, congestion, cough, diarrhea, fatigue, headache, loss of smell, runny nose, shortness of breath, sneezing, trouble breathing and weakness). Similarly, in the COVID recovered subgroup, one donor had experienced mild symptoms (fatigue, headache, loss of smell and taste) and one donor had experienced a more moderate symptomatology (body or muscle aches, cough, diarrhea, fatigue, fever, headache, loss of smell and taste and shortness of breath). Summaries of the demographics and SARS-CoV-2 serology test results are presented in Supplementary Table 2.

### Preparation of fixed blood clot samples

Healthy and COVID-19 fixed blood clot samples were commercially obtained (ZenBio) from a single donor, respectively. Whole blood was collected via venipuncture, added to an empty tube, and allowed to clot. The clot was mechanically disrupted and passed through a 30 μm filter in order to generate smaller clot fragments. Finally, the clots were fixed in 4% paraformaldehyde (PFA) and they were provided as 1 mL aliquots and stored at 4°C. For DHTM imaging, 200 μL of blood clots in aqueous solution was transferred in a 35-mm Ibidi ibiTreat μ-Dish (Ibidi GmbH, Germany) for DHTM imaging.

### Preparation of platelet-rich plasma

Platelet-rich plasma (PRP) was commercially obtained (BioIVT) for healthy and convalescent COVID-19 donors. Whole blood was collected in K2EDTA vacutainers and centrifuged in order to separate plasma from the cell pallet. PRP samples were transported and stored at -20°C. For all PRP samples, 50 μL of PRP was diluted in 200 μL of Alsever’s solution (Sigma-Aldrich, A3551) and 250 μL was transferred in a 35-mm Ibidi ibiTreat μ-Dish (Ibidi GmbH, Germany) for DHTM imaging.

### Label-free digital holo-tomographic microscopy

Label-free holo-tomographic imaging was performed using a 3D Cell Explorer microscope (Nanolive SA, Switzerland). Before each measurement, the petri dish containing either the fixed blood clots in aqueous solution or the plasma solution was placed in the microscope sample holder, and blood clots were allowed to sediment to the bottom of the petri dish for 10 minutes before imaging. DHTM was operated under standard laboratory conditions.

### Image processing and analysis

3D RI stacks obtained by DHTM were exported as TIFF files and imported into the open-source software Tomviz for 3D RI visualization. For the visualization of clot-specific structures, 3D stacks obtained by DHTM were digitally stained based on the RI values using STEVE (Nanolive SA, Switzerland), and each channel was exported in the form of a 3D stack as a TIFF file. The single-channel 3D stacks were imported into the open-source software Fiji and a 2x2x2 mean filter was applied as a noise removal filter. The single channels were merged into one image, exported as a TIFF file, and imported into Imaris 9.8 (Bitplane AG, Switzerland) in order to achieve a 3D surface segmentation. First, stacks were cropped along the *x*-axis and *y*-axis in order to exclude potential signal noise artefacts from the imaging process. Next, a surface was fitted for each channel with absolute intensity and automatic thresholding in order to achieve accurate signal segmentation. For the structural analysis and quantification of the blood clots, 3D RI stacks obtained by DHTM were imported into Imaris 9.8 and were cropped along the *x*-axis and *y*-axis in order to exclude potential signal noise artefacts. A 3x3x3 median filter was applied as a noise removal filter and a surface was fitted with absolute intensity and automatic thresholding. The morphologically relevant features were quantitatively measured, including the clot length, width, surface area, volume, and mean RI (Supplementary Table 3). The dry mass was calculated from the mean RI value of each blood clot, obtained from the 3D RI tomograms, using the following formula(*46*):

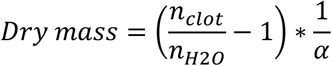

where *n*_*clot*_ is the mean RI value of the blood clot, *n*_*H*2*O*_ is the RI of water (1.333) and α is the wave-length-dependent RI increment, which was set to 0.001983 for λ = 520 nm(*47*).

### Raman spectroscopy setup and measurements

Raman spectra were obtained using NT-MDT NTEGRA Spectra system equipped with Olympus LMPLFLN 100× objective with numerical aperture NA = 0.8, and spectrometer grating of 600 g/mm. Samples were illuminated with the 561 nm wavelength laser (10 mW laser power at the sample). Spectra were acquired in the reflection mode, with an integration time of 20 s and 10 accumulations. To obtain a representative spectrum of the measured sample, multiple spectra were acquired in different locations of the sample and then averaged. Raman spectroscopy was conducted only on air-dried fixed blood clots in an aqueous solution deposited on SuperFrost glass slides.

## Acknowledgments

T.B. and P.N.N. thank Alex Dommann for strategic support. This work used the computational resources provided by the University of Bern, Switzerland. T.B. thanks Guillaume Witz at the Microscopy Imaging Center at the University of Bern for technical assistance. O.S. and P.N.N. thank Michel Calame for access to the Raman spectroscopy tool.

