## Supplementary Information for "Digital holo-tomographic 3D maps of COVID-19 microclots in blood to assess disease severity"

**This file includes:**

Supplementary Figure 1-6

Supplementary Table 1-3

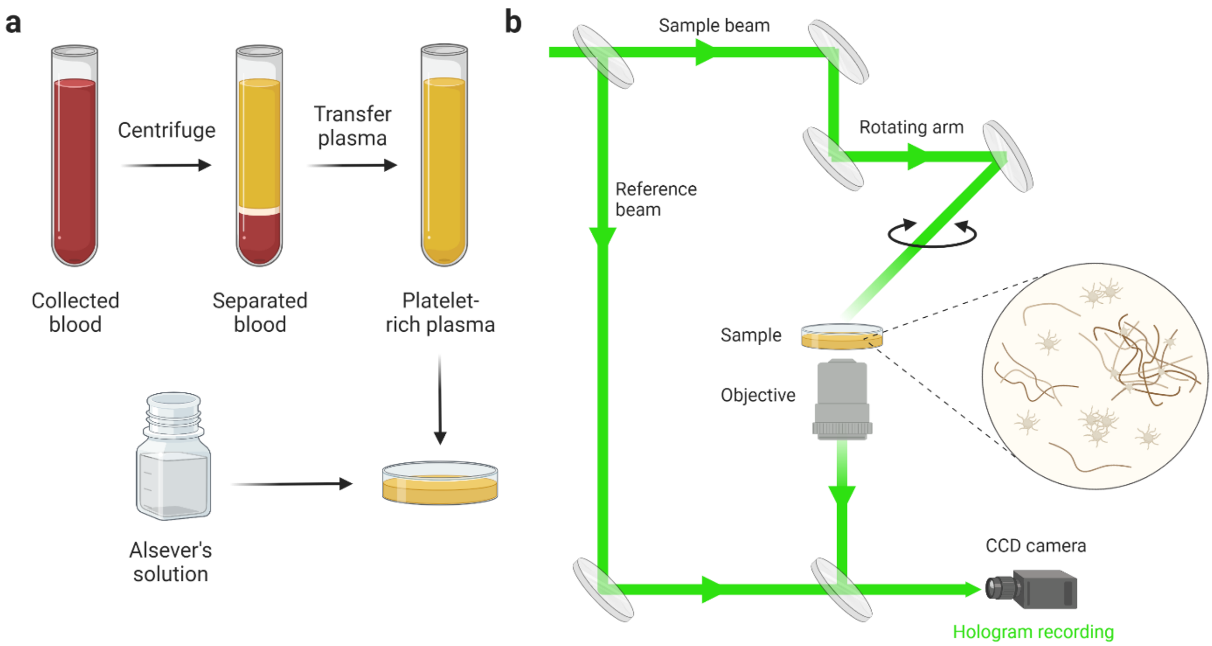


**Supplementary Figure 1: Sample preparation procedure and principle of DHTM for plasma imaging. a,** Platelet-rich plasma (PRP) was obtained after blood collection in K2EDTA vacutainers and centrifugation in order to separate each blood component. 50 µL of PRP is diluted in Alsever's solution at a final concentration of 1:5. 250 µL of plasma solution is transferred to a petri dish for imaging. **b,** Digital holotomographic microscopy uses a low-power laser beam ($\lambda$ = 520 nm) that splits into the reference and the sample laser beams before rejoining below the objective (60x, *NA* = 0.8), where the interference (hologram) is recorded. A rotational arm, located above the sample holder, rotates 360° around the sample at a 45° angle in order to obtain a 3D refractive index (RI) map. The presence of microclots and platelet pathology can be detected in a label-free manner from the 3D RI tomograms. Created with BioRender.com.

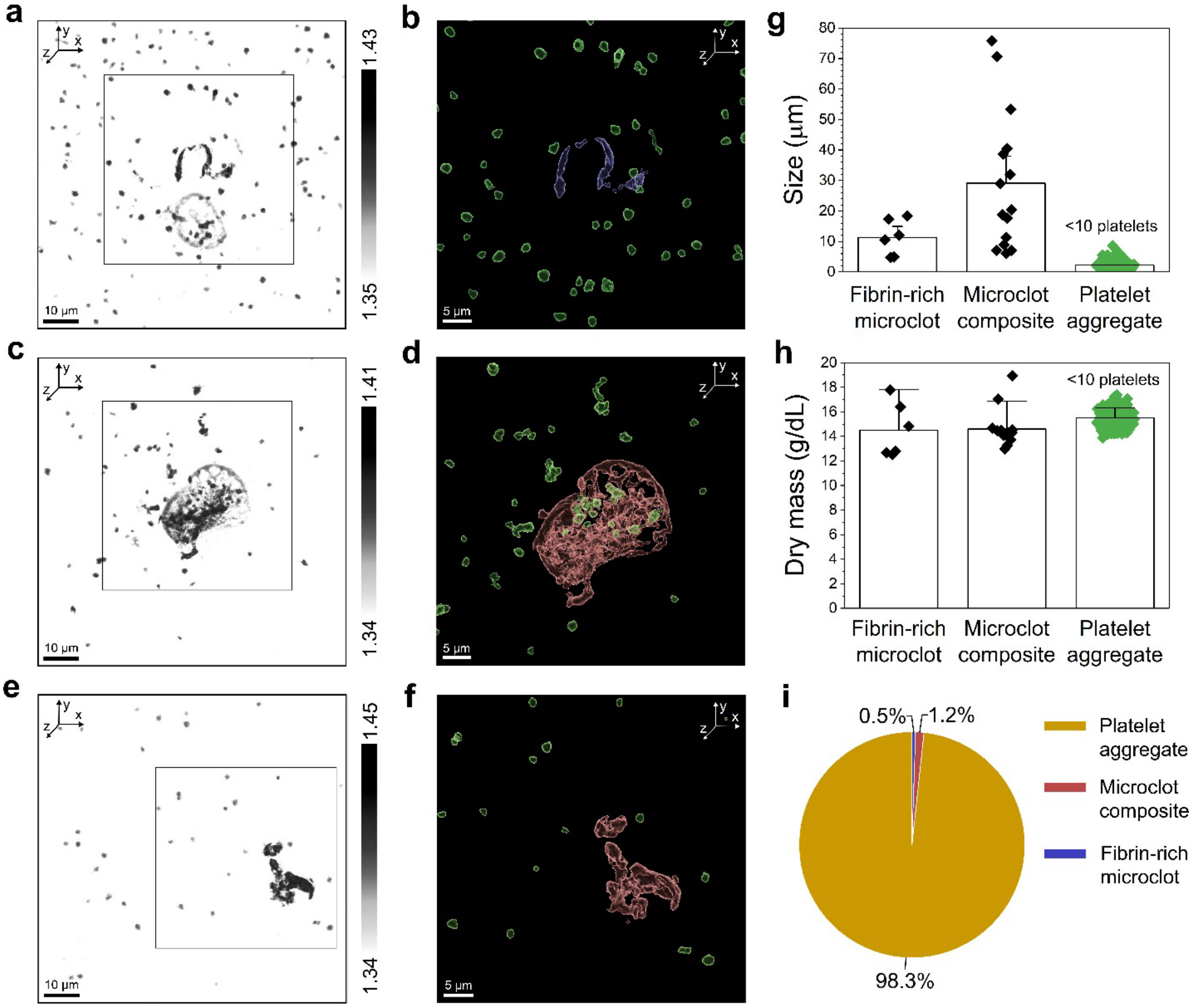


**Supplementary Figure 2: Structural analysis of microclots in platelet-rich plasma (PRP) of a COVID-19 positive donor without clinical symptomatology (PRP-CVDP3). a,** Refractive index (RI) tomogram of a fibrin-rich microclot in PRP. **b,** Corresponding segmented RI tomogram of the fibrin-rich microclot from the inset in **a**. **c,** RI tomogram of a microclot composite and platelet aggregates in PRP. **d,** Corresponding segmented RI tomogram of the microclot composite and platelet aggregates from the inset in **c**. **e,** RI tomogram of a microclot composite in PRP. **f,** Corresponding segmented RI tomogram of the microclot composite from the inset in **e**. **g,h,** Size and dry mass of fibrin-rich microclots, microclot composite structures and platelet aggregates (<10 platelets) in PRP. Error bars represent the standard deviation from the mean. **i,** Pie chart showing the microclot composition in PRP of donor PRP-CVDP3.

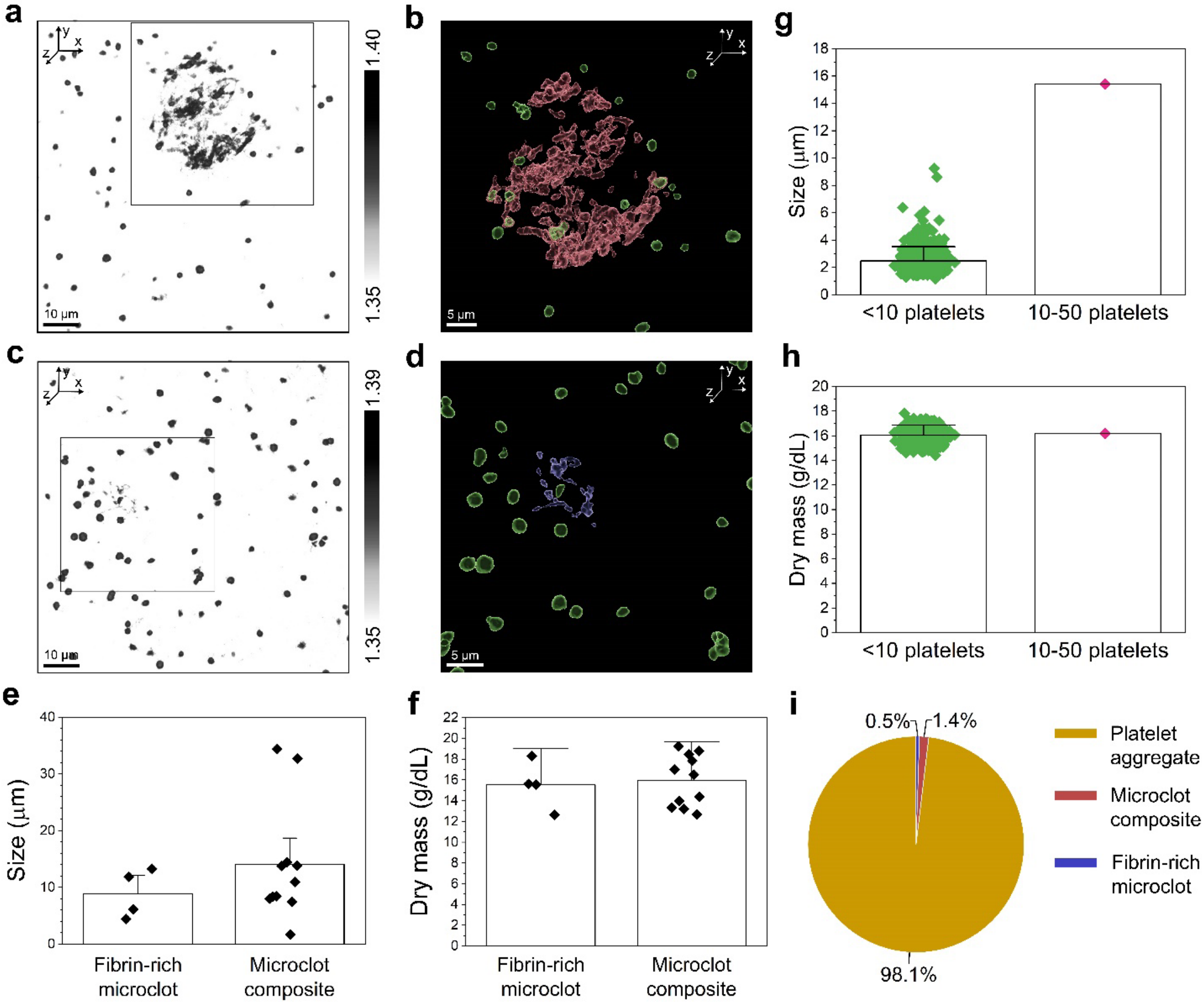


**Supplementary Figure 3: Structural analysis of microclots in platelet-rich plasma (PRP) of a COVID-19 recovered donor with moderate symptomatology (PRP-CVDR1). a,** Refractive index (RI) tomogram of a microclot composite and platelet aggregates in PRP. **b,** Corresponding segmented RI tomogram of the microclot composite and platelet aggregates from the inset in **a**. **c,** RI tomogram of a fibrin-rich microclot and platelet aggregates in PRP. **d,** Corresponding segmented RI tomogram of the fibrin-rich microclot and platelet aggregates from the inset in **c**. **e,f,** Size and dry mass of fibrin-rich microclots and microclot composite structures in PRP. **g,h,** Size and dry mass of platelet aggregates in PRP. Error bars represent the standard deviation from the mean. **i,** Pie chart showing the microclot composition in PRP of donor PRP-CVDR1.

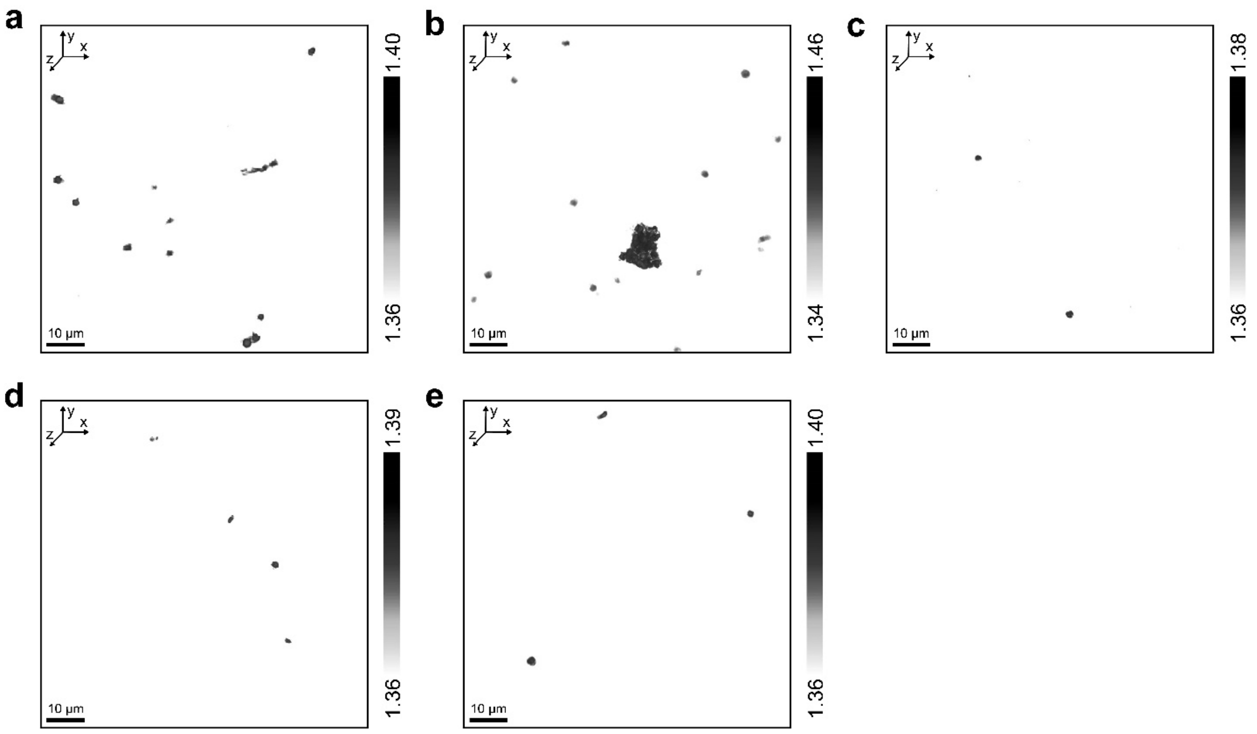


**Supplementary Figure 4: DHTM tomograms of platelet-rich plasma (PRP) of healthy controls (PRP-HC). a,** Refractive index (RI) tomogram of a small fibrin-rich microclot in PRP-HC1. **b,** RI tomogram of a microclot composite in PRP-HC2. **c,** RI tomogram showing individual platelets in PRP-HC3. **d,** RI tomogram showing individual platelets in PRP-HC4. **e,** RI tomogram showing individual platelets in PRP-HC5.

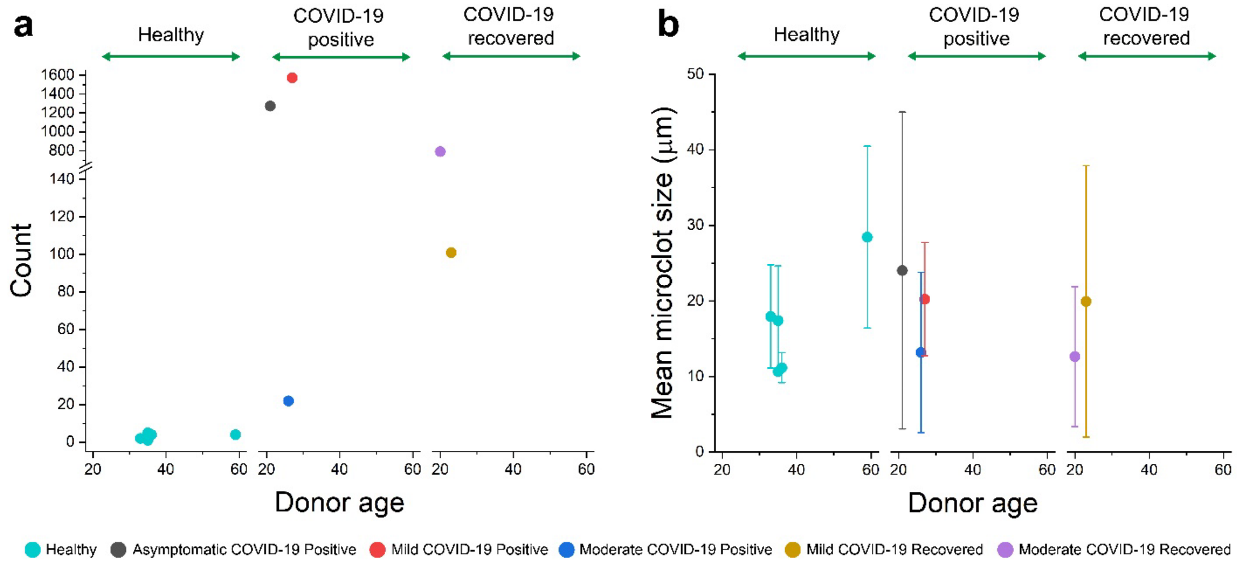


**Supplementary Figure 5: Age dependence of microclot prevalence and morphology in healthy and COVID-19 positive and recovered donors. a,** Prevalence of microclots detected in 75 µL of PRP from healthy and COVID-19 positive and recovered donors plotted against the donor age and grouped by healthy and COVID-19 subphenotypes. **b,** Mean microclot size of fibrin-rich microclots and microclot composites plotted against the donor age and grouped by healthy and COVID-19 subphenotypes.

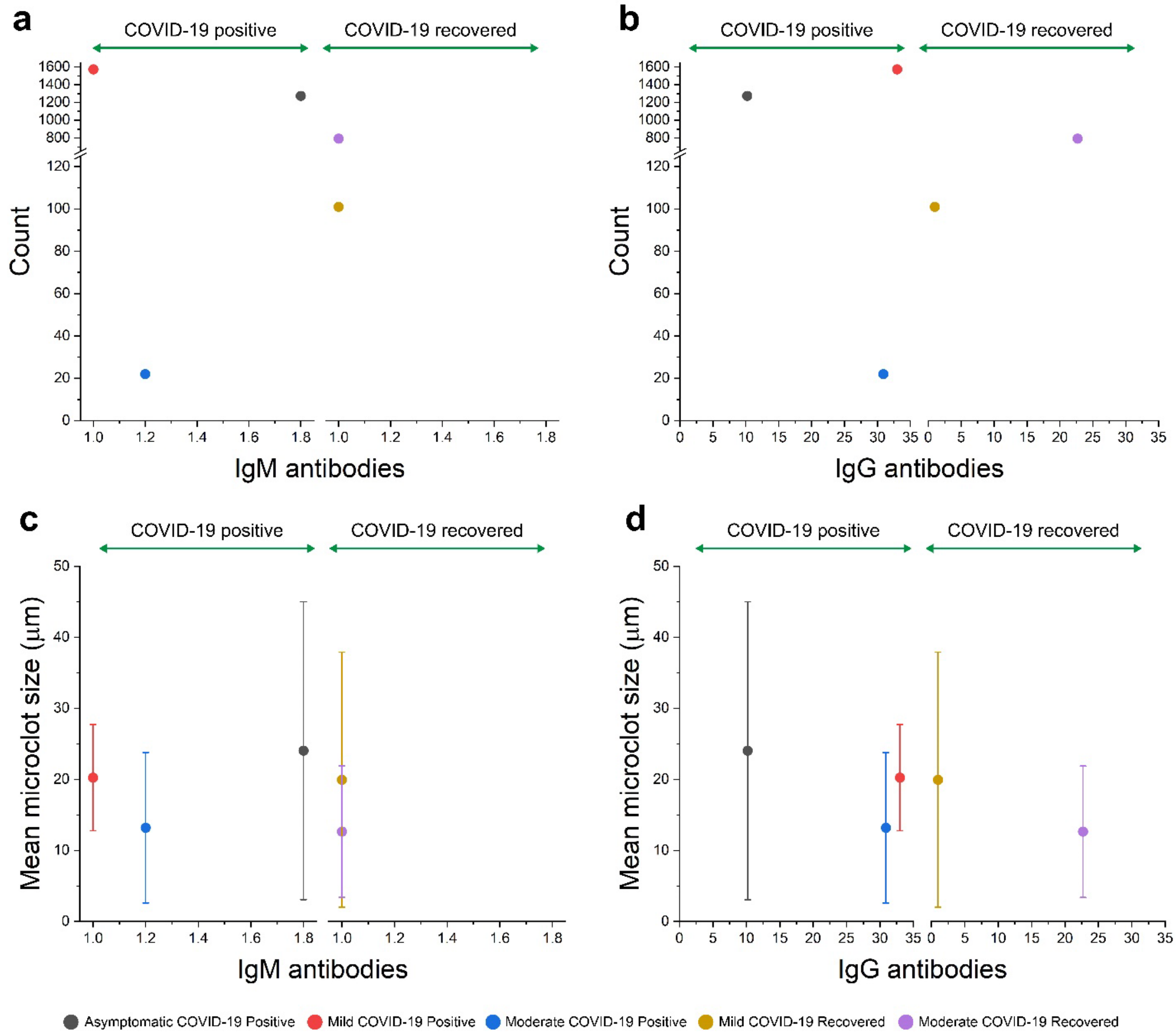


**Supplementary Figure 6: IgM and IgG antibodies dependence of microclot prevalence and morphology in COVID-19 positive and recovered donors. a,** Prevalence of microclots detected in 75 µL of PRP from COVID-19 positive and recovered donors plotted against the IgM antibodies level and grouped by COVID-19 subphenotypes. **b,** Prevalence of microclots detected in 75 µL of PRP from COVID-19 positive and recovered donors plotted against the IgG antibodies level and grouped by COVID-19 subphenotypes. **c,** Mean microclot size of fibrin-rich microclots and microclot composites plotted against the IgM antibodies level and grouped by COVID-19 subphenotypes. **d,** Mean microclot size of fibrin-rich microclots and microclot composites plotted against the IgG antibodies level and grouped by COVID-19 subphenotypes.

**Supplementary Table 1:** Summary of the demographics for the samples of the fixed blood clots in aqueous solution.

| | Fixed blood clots | |
|---|---|---|
| | **FC-HC** | **FC-CVD** |
| **Donor classification** | Healthy | COVID-19 convalescent |
| **Gender** | F | F |
| **Age** | 49 | 42 |
| **Ethnicity** | African American | African American |
| **Collection date** | - | 15/09/2021 |

**Supplementary Table 2:** Summary of the demographics, SARS-coV-2 serology test results (IgM and IgG antibodies), symptomatology and sample collection details for healthy and COVID-19 platelet-rich plasma (PRP) samples.

| | Platelet-rich plasma (PRP) | | | | | | | | | |
|---|---|---|---|---|---|---|---|---|---|---|
| | **PRP-HC1** | **PRP-HC2** | **PRP-HC3** | **PRP-HC4** | **PRP-HC5** | **PRP-CVDP1** | **PRP-CVDP2** | **PRP-CVDP3** | **PRP-CVDR1** | **PRP-CVDR2** |
| **Donor classification** | Healthy | Healthy | Healthy | Healthy | Healthy | COVID-19 positive | COVID-19 positive | COVID-19 positive | COVID-19 recovered | COVID-19 recovered |
| **Gender** | F | F | F | M | M | F | F | F | F | F |
| **Age** | 35 | 33 | 59 | 36 | 35 | 26 | 27 | 21 | 20 | 23 |
| **Ethnicity** | White | White | Other | White | Asian | Hispanic/ Latino | White | White | White | White |
| **SARS-coV-2 serological test** | | | | | | | | | | |
| IgM antibodies | - | - | - | - | - | Positive (1.2) | Negative (1) | Positive (1.8) | Negative (1) | Negative (1) |
| IgG antibodies | - | - | - | - | - | Positive (30.9) | Positive (33) | Positive (10.2) | Positive (22.7) | Negative (1) |
| **Symptomatology** | - | - | - | - | - | Moderate | Mild | Asymptomatic | Moderate | Mild |
| **Diagnosis date** | - | - | - | - | - | 17/12/2020 | 07/11/2020 | 01/10/2020 | 12/10/2020 | 16/11/2020 |
| **Collection date** | 19/12/2022 | 04/07/2023 | 04/07/2023 | 04/07/2023 | 04/07/2023 | 13/01/2021 | 04/12/2020 | 20/10/2020 | 13/01/2021 | 06/01/2021 |
| **Days from diagnosis** | - | - | - | - | - | 27 | 27 | 19 | 93 | 51 |

**Supplementary Table 3:** Description of the quantified morphological parameters extracted from Imaris 9.9.

| Morphological parameter | Description |
|---|---|
| **Length/Size** | The length of the longest principal axis inside the object (*BoundingBoxOO Length C)* |
| **Width** | The length of the second longest principal axis inside the object (*BoundingBoxOO Length B)* |
| **Surface area** | The sum of the triangle surfaces |
| **Volume** | Quantification of how much a surface object occupies |
| **Mean RI** | Mean intensity of voxels enclosed within the surface |
